# No preference for prosocial ‘helping’ behavior in rats with concurrent social interaction opportunities

**DOI:** 10.1101/2020.11.18.388702

**Authors:** Kelsey A. Heslin, Michael F. Brown

**Author notes:** <u>Corresponding Author:</u> Kelsey Heslin. **Open Practices Statement** The datasets generated and analyzed during the current study are available from the corresponding author upon request. This study was not preregistered. Current address: University of Iowa, Department of Psychiatry, 500 Newton Road Iowa City, IA 52242, United States.

## Abstract

‘Helping behavior’ tasks are proposed to assess prosocial or ‘empathic’ behavior in rodents. This paradigm characterizes the behavior of subject animals presented with the opportunity to release a conspecific from a distressing situation. Previous studies found a preference in rats for releasing restrained or distressed conspecifics over other controls (e.g., empty restrainers or inanimate objects). An empathy account was offered to explain the observed behaviors, claiming subjects were motivated to reduce the distress of others based on a rodent homologue of empathy. An opposing account attributes all previous results to subjects seeking social-contact. To dissociate these two accounts for helping behavior, we presented subject rats with three simultaneous choice alternatives: releasing a restrained conspecific, engaging a non-restrained conspecific, or not socializing. Subjects showed an initial preference for socializing with the non-restrained conspecific, and no preference for helping. This result contradicts the empathy account, but is consistent with the social-contact account of helping behavior.

---

Social interaction paradigms are valuable tools for testing whether experimental factors influence social behaviors in animal models. These tasks may provide insights into abnormal social behavior seen in neurodevelopmental and neuropsychiatric disease (Silverman et al., 2010). Focus on animal models of abnormal social interactions has increased demand for a more diverse array of behavioral assays that capture disease-relevant aspects of social behavior (Bishop & Lahvis, 2011; Nestler & Hyman, 2010; Silverman et al., 2010; Stewart & Kalueff, 2015). The helping behavior paradigm is one such purported assessment, aimed at measuring prosocial behavior in rodents. However, this paradigm’s ability to evaluate prosocial behavior beyond the confound of social reward has not been demonstrated.

The helping behavior paradigm examines the behavior of an acting subject rat when given the opportunity to ‘rescue’ or help a distressed conspecific (Bartal et al., 2011). This task was initially proposed as a method for exploring whether rodents are capable of displaying empathy toward conspecifics. Due to philosophical and operational complexities of the term empathy, some researchers have suggested the helping behavior captured by Bartal et al. (2011) may better be described as prosocial behavior (Hollis & Nowbahari, 2013; Vasconcelos et al., 2012). Prosocial behavior is any behavior that provides a benefit to another individual, with little or no cost to the actor (Schwab et al., 2012; Vasconcelos et al., 2012). The helping behavior paradigm has since been proposed as an assessment for prosocial behavior in rodent models, aiming to identify the etiology of disordered prosocial behaviors (Cox & Reichel, 2020; Meyza & Knapska, 2018; Millan & Bales, 2013; Mony et al., 2018; Wöhr & Scattoni, 2013). However, the methods used in existing studies have not directly ruled out the alternative possibility that subjects in the task behave to obtain social reward, rather than help. The objective of the current study was to provide data needed to evaluate which of these explanations is more likely.

Generally, the helping behavior paradigm examines the reaction of a freely moving subject rat in an arena, in response to a presumably distressed stimulus rat. In most cases, stimulus rat distress is produced by means of confinement to a physical restrainer (e.g., a restraining tube; Bartal et al., 2011; Bartal et al., 2014; Havlik et al., 2020; Silberberg et al., 2014). Restraining tubes are commonly used to induce distress in laboratory rodents without causing physical pain or injury (Sun et al., 2013; Ward, 1972; Ward et al., 2003; Zimprich et al., 2014). In the original Bartal et al. (2011) study, subject rats were observed approaching and releasing restrained conspecifics by physically manipulating the weighted door of the restrainer (Bartal et al., 2011). Importantly, opening the restrainer door released the trapped stimulus rat into the same area as the subject rat, thereby allowing social interaction. The door-opening behavior, (i.e., the prosocial behavior of interest) was compared in terms of frequency and latency across separate conditions. Initially, subjects were more likely to perform door-opening when the restrainer contained a cagemate compared to an inanimate object or an empty restrainer. Additionally, subjects performed door-opening behavior at increasingly shorter latencies as the 12 days of testing progressed in the trapped cagemate condition, but not in the compared empty and object control conditions (Bartal et al., 2011).

Following this phase, Bartal and colleagues (2011) carried out additional testing of subjects previously exposed to the trapped cagemate condition to compare door-opening behavior in an additional ‘separated’ condition. In this case, restrained cagemates were now released into a distal chamber, preventing social contact from reinforcing subjects’ door-opening for 27 additional days. Subjects continued to release restrained cagemates even when social contact was no longer available as a reward for door-opening. From this, the authors concluded that subjects were likely acting based on a rodent homologue of empathy (Bartal et al., 2011).

In response to this empathy-based conclusion, Silberberg and colleagues (2014) performed a slightly altered replication of Bartal et al. (2011) to examine an alternative explanation for the observed results. Silberberg et al. (2014) argued that the control conditions used by the original study could not adequately rule out the hypothesis that the observed prosocial behavior resulted from subjects acting to gain social contact with the restrained stimulus rats. This consideration is warranted, as social contact can act as a powerful reinforcer for rats (Evans et al., 1994; Hiura et al., 2018). To address this potential confound, Silberberg and colleagues (2014) demonstrated that when the same conditions were presented to subjects in a different order, (i.e., separated, then trapped, then separated again) subject behavior did not support an empathy-based interpretation. That is, when subjects were first exposed to the ‘separated’ condition instead of the socially rewarding ‘trapped’ cagemate condition, door-opening latencies increased, and door-opening frequency decreased over time. Door-opening latency only decreased when subjects were newly exposed to the trapped condition in phase 2 of the study (Silberberg et al., 2014). Furthermore, once exposed to social reward for door opening, Silberberg et al. found that door-opening behavior did not extinguish, and latencies did not increase when subjects were switched back to a separated condition with no social contact reward for 27 additional sessions (Silberberg et al., 2014). Therefore, enduring expectation of social contact, here called the social-contact hypothesis, may better explain the pattern of results reported by both Silberberg et al., (Schwartz et al., 2017; Silberberg et al., 2014) and Bartal et al. (2011).

While this follow-up approach illustrated the need to consider a social contact explanation for helping behavior, it did not directly pit the two hypotheses against each other methodologically. Subsequent studies have further explored the role of factors like subject familiarity, additional observers, and anxiolytic treatment in the helping behavior paradigm but in almost all cases, never offer simultaneous social interaction opportunities to directly assess the confound of social contact motivation (Bartal et al., 2014, 2016; Carvalheiro et al., 2019; Havlik et al., 2020; Sato et al., 2015). Comparing helping behavior frequency and latency between experimental and control conditions does not directly reveal subject preference. One study that did directly assess behavior using concurrent social options found that subject rats choosing between approaching a non-restrained rat or approaching and releasing a restrained rat chose these two options at equal frequency (Hachiga et al., 2018). In two additional conditions, subjects preferred the restrained rat option over an empty goal box option and the non-restrained rat option over the empty goal box (Hachiga et al., 2018). This preference for any social option but no preference for helping the restrained rat supported the social-contact hypothesis. However, this approach only compared subject preference between two of the three stimulus options at any given time.

Before the validity of the helping behavior paradigm can be established, the social contact explanation must be thoroughly assessed. If subjects are behaving to obtain rewarding social contact, this undermines claims of empathic or prosocial motivations. To parse these competing interpretations, we modified this helping behavior paradigm to allow the dissociation of these two explanatory accounts. We presented subject rats with three simultaneous choices: releasing a restrained conspecific followed by social contact, socially engaging a non-restrained conspecific, or not socializing. These three options were made distinct by spatial location, but all required equal effort to access. The social-contact hypothesis and the empathy hypothesis of helping behavior make diverging predictions in this three-choice scenario. Subjects preferring locations containing a rat (i.e., a social contact opportunity), but no preference between restrained or non-restrained stimulus rats would support the social-contact hypothesis. Non-preferential results for the two social options would be consistent with the interpretation that prosocial helping behavior is not produced by altruism or empathy, but rather by a motivation for social contact, regardless of the conspecific’s distress state. In contrast, a preference for releasing the restrained animal only would support the empathy hypothesis.

## Method

### Subjects

Twenty male Sprague-Dawley subject rats, age 9 months (Envigo, Indianapolis, IN), chose between releasing a restrained conspecific, engaging with a non-restrained conspecific, or avoiding social interaction in 24 total choice trials. One subject was excluded from analyses for failing to make a choice in 66% of test sessions, resulting in 19 subjects total. A separate cohort of 24 Sprague-Dawley males (age 4 months; Envigo) served exclusively as stimulus rats. All rats were housed in pairs on a reverse 12h light/dark cycle with unlimited access to food and water. All experimental procedures were approved by the Villanova University Institutional Animal Care and Use Committee and followed applicable international, national, and institutional guidelines for the care and use of animals.

### Apparatus

Pre-training and testing took place in a custom 1.2 m square arena (Figure 1a) with 14 cm flat black walls, containing three identical transparent choice chambers. The choice chambers (45 cm × 24 cm × 20 cm) were identical in size to the home cages of the animals and were composed of perforated polycarbonate. Chambers had twenty-six 0.32 cm holes distributed across the four sides to make olfactory and auditory cues within the chamber available to subjects outside. Each choice chamber was accessible by a single inward-swinging door that subjects could push open from outside (Figure 1b). The circular entry hole was 7.6 cm in diameter and covered by a rectangular, transparent 10 × 12.5 cm polystyrene door on the interior of the cage, secured from the top with a metal hinge. An internal perforated divider (polystyrene grid with 1.2 × 1.2 cm square cells) divided each choice chamber in half to separate stimulus rats from the entry door. The divider prevented non-restrained stimulus rats from moving closer to the entry door than restrained rats, and prevented subject rats from entering the restrainer, but still allowed for social contact through the perforations.

**Figure 1.**
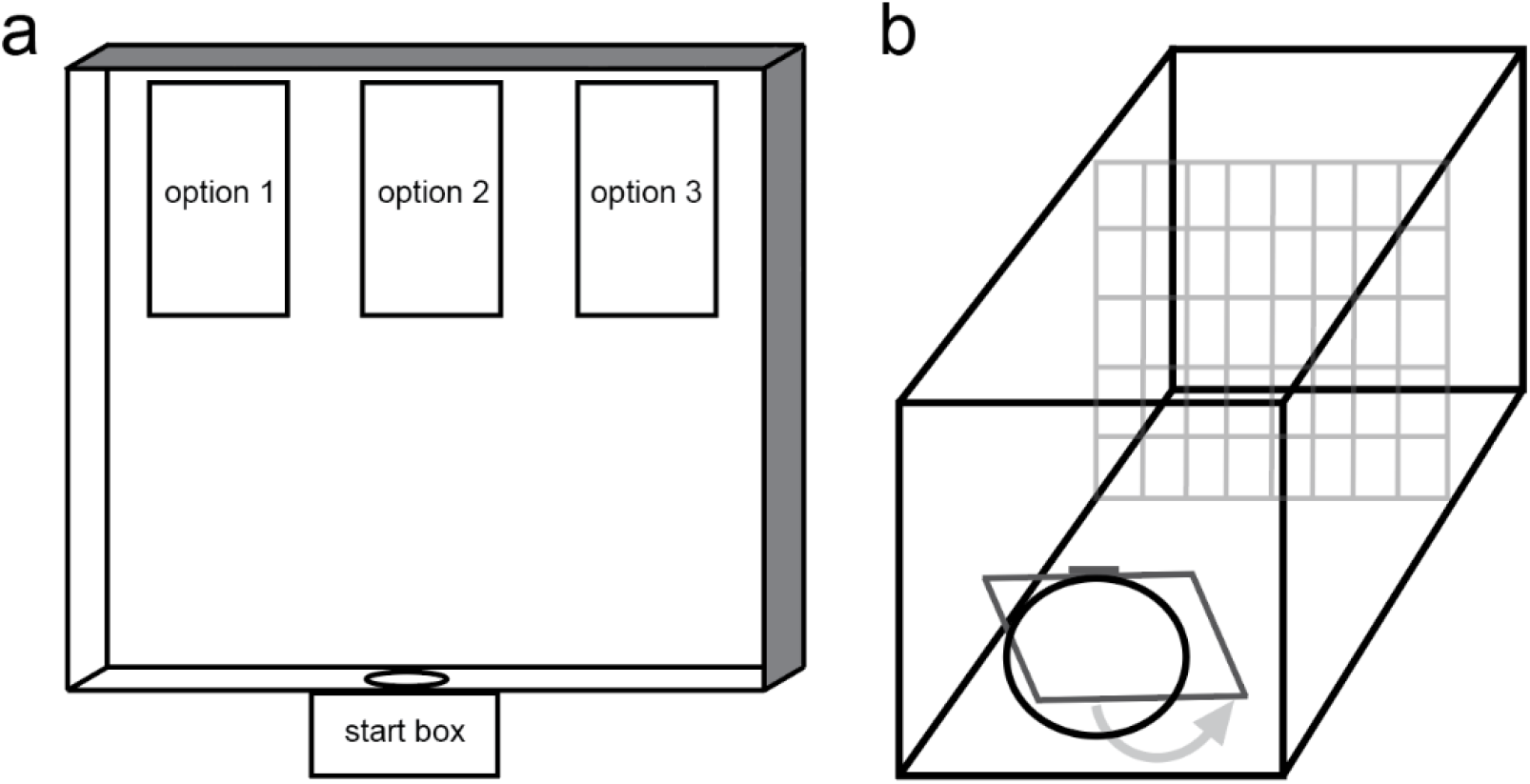
(a) Diagram of the task arena (top view). All three choice chambers options were spatially counterbalanced by trial. Subjects always entered the arena from the start box. (b) Choice chamber diagram. Subjects were able to enter each choice chamber by applying pressure to a transparent, in-ward swinging door. Once inside any given chamber, subjects could interact with stimulus animals located on the other side of the perforated divider (shown in grey). The presence of the divider limited the range of motion for the non-restrained stimulus rats.

### Pre-training

Subjects were habituated to the task arena and trained in chamber door operation prior to testing. On pre-training day 1 (PT1), subjects explored the arena with no additional rats or food present for 10 minutes. On PT2-PT4, subjects could explore the arena and feed from containers holding palatable sucrose pellets. Feeding containers were placed on the side of the arena where the choice chambers would subsequently be located, in each of the three possible locations. To avoid spatial preferences, once a subject fed at one location, that food container was removed for the following trials. For PT5-7, the choice chambers were added to the arena with sucrose pellet containers inside all three choice chambers. No doors covered the entrances to the choice chambers at this stage in training. All chambers contained sucrose on PT5. If a subject fed at one location, that location was not available in successive trials until the two remaining locations were chosen. On all following PT days (PT8-13), the sucrose pellet containers were again placed inside all three choice chambers, but subjects were required to push open doors in order to access the sucrose. This training phase was repeated until each subject displayed door-opening behavior for three consecutive days. Visited locations were removed until all remaining locations were visited on subsequent trials. Subjects required between 10 and 13 days to complete pre-training, with a mean of 11.2 days to meet criteria.

### Testing

Testing sessions were broken into two blocks of 12 sessions each to match the initial number of testing sessions used by Bartal et al. (2011) and to also assess whether experience in the task influenced choice behavior. The 24 total choice sessions occurred once daily over 24 days. For every choice trial, each of the three spatially counter-balanced choice chambers contained either a stimulus rat in a transparent polycarbonate perforated restraining tube (Thomas Scientific), a stimulus rat not in a restraining tube, or no stimulus rat. To maintain an equal level of familiarity, stimulus rats were used on a rotating schedule resulting in an equal number of exposures for each subject rat. For every choice trial, subjects entered the arena from a start box via a guillotine door. To avoid any effects of location biases, the spatial location of each of the three stimulus options was counterbalanced across trials for all subjects. All choice trials lasted until a choice was made by the subject rat, or until eight minutes in the arena expired. A choice was defined as the subject rat opening the door and entering the choice chamber with all four paws inside. Only one choice per trial was allowed. Entering the choice chamber containing the restrained rat resulted in immediate release of the restrainer door, allowing the restrained rat to escape within the choice chamber. Therefore, choosing to enter either chamber containing the restrained or non-restrained stimulus rat resulted in social contact through the perforated divider. No additional physical effort was required to release the restrained rat. To ensured that the duration of social reward was equal for both the restrained and non-restrained rat options, subject rats remained in the chosen chamber for one minute after entering, regardless of chamber type chosen.

### Analyses

The planned analyses were aimed to dissociate competing predictions made by the empathy and social-contact hypotheses. As choice data were collected in the form of proportions derived from frequency counts, these data were transformed to arcsine values for analysis using the following equation:

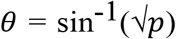

Transformed choice proportions for each subject were used for computing five separate *t*-tests and corresponding Bayes factors. Comparison values for chance were transformed identically. All Bayes factors were calculated with JZS priors using the method described by Rouder et al. (Rouder et al., 2009). All t-tests used the arcsine value of choice proportions per subject within block. A significance level of α =.05 was used for all statistical analyses.

To assess any effects of choice type or block on choice latencies, we used a 3 × 2 (Choice Type × Block) repeated measures ANOVA with individual subject mean latencies. One subject did not make any choices to the ‘empty’ option in Block 2. This missing value was replaced using mean substitution because it constituted less than 1% of the data and because our hypotheses do not depend on latency variability for interpretation (Rubin et al., 2007).

## Results

Choice data from 24 total testing sessions were analyzed in two blocks of 12 in order to observe changes in choice patterns as subject familiarity with the task increased. Choices made by the 19 subjects indicated an overall preference for selecting the non-restrained rat chamber (Figure 2a). In Block 1, subjects chose the non-restrained rat location 47.56% of the time, the restrained rat location 29.78%, and the empty location 22.67%. Because both the empathy hypothesis and the social-contact hypothesis predict preferences for choosing to approach chambers containing rats, an initial set of t-tests determined if a preference emerged for choosing social choice chambers (i.e., chambers containing either the restrained or non-restrained rat) compared to chance (0.67). Subjects in Block 1 displayed a significant preference for making choices to social chambers *t*(18) = 4.174, *p* = 0.001, with a Bayes factor of 60.32 in favor of the alternative, reflecting a likelihood ratio of 60.32:1 in favor of the alternative hypothesis (i.e., that the sample mean is different than the comparison value; Rouder et al., 2009).

**Figure 2.**
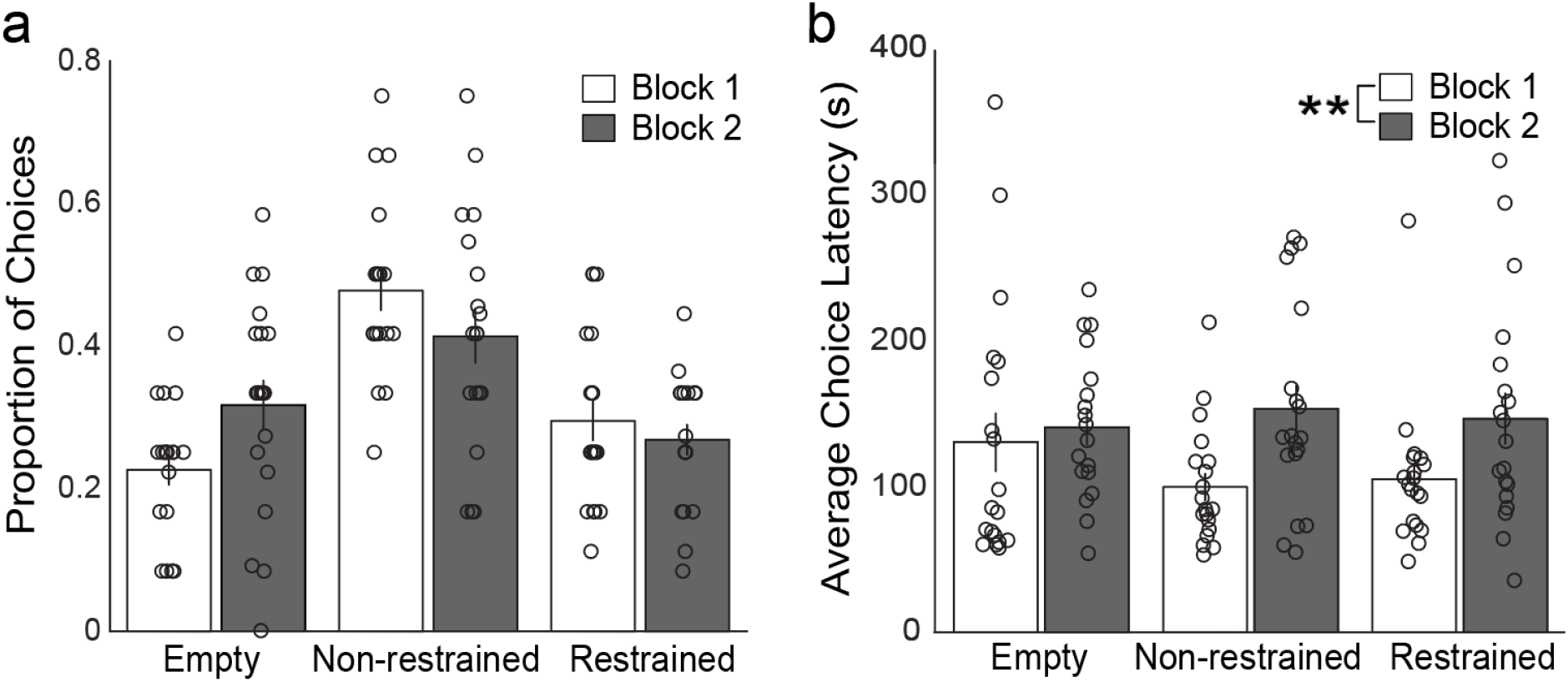
(a) Proportion of choices made to each stimulus option by trial block. Each open circle represents an individual subject’s proportion of choices to each stimulus chamber type in the first 12 choice sessions (Block 1) and the following 12 sessions (Block 2). Subjects preferred to choose the non-restricted stimulus at option in Block 1 but had no significant preference in Block 2. (b) Latency to chamber choice. Open circles represent individual subject mean latency (in seconds) to each stimulus chamber type divided between the first 12 choice sessions (Block 1) and the following 12 sessions (Block 2). Overall latency to make a choice was higher in the second block but did not vary by stimulus type. ** *p* value < 0.01; error bars reflect SEM.

In the second block of choice sessions, subjects again selected the non-restrained option most often with 41.36% of choices made to that location, 26.82% to the restrained location, and 31.82% of choices to the empty location. However, subjects in Block 2 no longer displayed a significant preference for choosing social chambers compared to chance, *t*(18) = 0.855, *p* = 0.404, with a Bayes factor of 3.05 in favor of the null.

To further evaluate the two competing hypotheses, choices to the restrained rat over the non-restrained rat were examined, as the empathy hypothesis would predict a significant preference for choosing to enter the chamber of the restrained animal, causing its release. To assess this, the proportion of choices to the restrained rat out of only the two social options was compared to chance (0.5). Subjects significantly preferred approaching the non-restrained over the restrained rat in both Block 1, *t*(18) = 3.629, *p* = 0.002, with a Bayes factor of 21.05 in favor of the alternative, and in Block 2, *t*(18) = 2.446, *p* = 0.025, with a Bayes factor of 2.458 in favor of the alternative. Additionally, a paired samples t-test showed no significant differences in choices to the restrained rat chamber as a function of block *t*(18) = 0.579, *p* = 0.570, with a Bayes factor of 3.624 in favor of the null. While no preference for releasing the restrained rat was observed when considering all subjects pooled together, we also examined whether individual subjects developed a preference for helping. In Block 1, only 3 out of the 19 subjects chose to release the restrained rat more frequently than choosing the other two stimulus options. In Block 2, 0 of the 19 subjects chose the restrained rat chamber more frequently than the other two options.

As shown in Figure 2b, latency to choose a stimulus chamber did not significantly vary based on the stimulus type chosen, but choice latencies did increase overall with more exposure to the task. A 3 × 2 (Choice Type × Block) repeated measures ANOVA indicated that choice latency was not significantly impacted by the type of chamber chosen *F*(2,36) = 0.384, *p* = 0.684 but did significantly increase in Block 2 *F*(1,18) = 9.444, *p* < 0.007, with no significant interaction *F*(2,36) = 1.933, *p* = 0.159.

## Discussion

To evaluate competing explanations for previously observed helping behavior, we tested whether subject rats chose between three concurrent options: releasing a restrained rat, engaging a non-restrained rat, or not socializing. Our findings indicate that subjects initially preferred choosing locations that resulted in socialization opportunities, particularly with non-restrained animals. This pattern of results does not support the empathy account posited by Bartal et al. (2011) because subjects did not prefer to release the restrained animal in either block of choice sessions. Instead, our data are more consistent with the social-contact hypothesis (Schwartz et al., 2017; Silberberg et al., 2014) and past studies showing that subjects do not prefer to release a restrained conspecific when given a concurrent choice of a non-restrained conspecific (Hachiga et al., 2018).

Our findings add to existing evidence which indicates that helping behavior is not the result of purely prosocial motivation when subjects are given multiple options. Instead, helping behavior may be the result of subjects acting to gain social contact or avoid aversive distress signals. For example, Hachiga et al. (2018) showed that subject rats to do not prefer to release a restrained conspecific when given a second option to interact with a non-restrained rat. Instead, subjects chose both social options with similar frequency, suggesting that their selections were driven by the resulting social contact opportunity, and not by relieving the distress of the restrained rat. Furthermore, rats given the option to either escape the testing arena or help a restrained conspecific engage in less helping behavior and do so at longer latencies than subjects that do not have the option to escape (Carvalheiro et al., 2019). This finding contradicts the empathy hypothesis which would predict equal helping frequencies and latencies regardless of the ability to escape. Instead, these results suggest that avoiding either the bright arena or the distress of the conspecific is more motivating than relieving that distress. Another study using a progressive ratio operant schedule showed that subjects were less motivated to perform the operant helping behavior when the behavior did not result in social interaction compared to a condition when it did result in socializing (Cox & Reichel, 2020). In another version of the helping behavior paradigm where discomfort was induced with water rather than restraint (Sato et al., 2015), follow-up studies demonstrate that the reported helping behavior can be attributed to the reinforcing properties of social contact and water (Schwartz et al., 2017). As whole, these findings cast doubt on the interpretation that the helping behavior paradigm is an assessment for prosocial or ‘empathic’ behavior in rodents. Rather, the observed behavior of subjects can be explained by seeking reinforcement independent of relieving distress. Our findings add to this body of work by directly showing that subjects do not prefer to help restrained rats when other choice options are present. Rather, subjects initially preferred to socialize with a non-restrained stimulus rat and did not display any preference for helping the restrained rat.

While our findings support the social-contact hypothesis, our pattern of results differ slightly from similar studies. For example, Hachiga et al. (2018) found that when offered two choices of visiting a restrained or non-restrained conspecific, subjects showed no preference between these two social options. Instead, we observed that when three options are available, subjects initially preferred the non-restrained stimulus animal over the restrained one. Our subjects may have preferred to avoid the non-restrained animals if auditory or olfactory distress signals from the restrained animals were perceived to be aversive and elicited an avoidance response (Hollis & Nowbahari, 2013; Rogers-Carter et al., 2018; Vasconcelos et al., 2012). However, we did not measure ultrasonic vocalizations (Borta et al., 2006; Knutson et al., 2002) to confirm that distress calls impacted subject behavior. Additionally, selection of the restrained rat location may have resulted in a slightly longer delay to social contact compared to the non-restrained animal, as a short amount of time was necessary to exit the restraining tube upon door release. More stringent control over the delay period between location selection and social contact onset across options may yield different results. In any case, the present choice results for Block 1 are consistent with the social-contact hypothesis and offer no support to the empathy hypothesis.

Moreover, latency to location choice was not affected by the stimulus type chosen, revealing no particular urgency to release the restrained rat. Choice latency increased overall from the first block to the second block of test sessions. The lack of a significant preference for social choices and the overall choice latency increase in Block 2 may reflect decreased social motivation over time as subjects learned that only limited physical interaction was available through the internal chamber divider for a limited amount of time (one minute per trial). Providing a social encounter of greater value may have prevented this late decline in choices to social contact opportunity locations.

A limitation of note is that we did not use the cagemates of subject rats as stimulus animals, as our design required two concurrent stimulus animals, a demand that was not compatible with our animal housing facility practices. Our use of unfamiliar stimulus rats diverged from past studies that specifically used cagemates as stimulus animals (Bartal et al., 2011; Silberberg et al., 2014). Familiarity with the restrained stimulus rat may modulate the reaction of subjects, with familiarity increasing prosocial behaviors in some instances (Ben-Ami Bartal et al., 2014; Rogers-Carter et al., 2018). Another limitation to consider is our use of exclusively male subjects. Some past reports indicate a possible sex difference in prosocial behavior frequency (Bartal et al., 2011; Rogers-Carter et al., 2018). For example, Bartal et al. (2011) noted that female subjects were more likely to learn to perform helping behavior and to do so at shorter latencies. Using exclusively male rats in this experiment may have prevented the emergence of preference for approaching the restrained rat. However, Silberberg and colleagues’ exclusive use of female rats did not support the claim of females acting empathically (Silberberg et al., 2014).

In general, our findings support the social-contact hypothesis as a viable interpretation of helping behavior, indicating that motivation for social reward plays a nontrivial role in the observed outcomes. Consequently, the inclusion of control conditions that account for this critical confound are needed for all variations of the helping behavior paradigm. Existing helping behavior findings should be interpreted cautiously, especially regarding translational applications to human empathic behavior.

## Notes

**Author Note** Data from this experiment served as a portion of K.H.’s Master’s thesis at Villanova University. The authors declare no conflicts of interest.

### Competing Interest Statement

The authors have declared no competing interest.

